# Alternative splicing analysis benchmark with DICAST

**DOI:** 10.1101/2022.01.05.475067

**Authors:** Amit Fenn, Olga Tsoy, Tim Faro, Fanny Rössler, Alexander Dietrich, Johannes Kersting, Zakaria Louadi, Chit Tong Lio, Uwe Völker, Jan Baumbach, Tim Kacprowski, Markus List

## Abstract

Alternative splicing is a major contributor to transcriptome and proteome diversity in health and disease. A plethora of tools have been developed for studying alternative splicing in RNA-seq data. Previous benchmarks focused on isoform quantification and mapping. They neglected event detection tools, which arguably provide the most detailed insights into the alternative splicing process. DICAST offers a modular and extensible framework for the analysis of alternative splicing integrating 11 splice-aware mapping and eight event detection tools. We benchmark all tools extensively on simulated as well as whole blood RNA-seq data. STAR and HISAT2 demonstrated the best balance between performance and run time. The performance of event detection tools varies widely with no tool outperforming all others. DICAST allows researchers to employ a consensus approach to consider the most successful tools jointly for robust event detection. Furthermore, we propose the first reporting standard to unify existing formats and to guide future tool development.

## Introduction

Alternative splicing (AS) affects around 95% of eukaryotic genes with multiple exons^1,2^ and gives rise to a large number of isoforms. AS is involved in cellular processes and disease development (see recent reviews on cancer^3^, muscle^4^, and neuron^5^ development). The most popular technology to study AS is short-read RNA sequencing (RNA-Seq). Possibilities for AS analysis from short-read RNA-Seq data comprise splice-aware mapping, *de novo* transcriptome assembly, AS detection and/or quantification on the exon, isoform, or event level as well as differential splicing analysis. Each year, several new tools are published for each of these analysis types.

Existing benchmark studies that could guide users on which tool to use for which analysis have several limitations. First, the studies use as ground truth either only a small subset of well-studied genes or simulated RNA-Seq data with randomly introduced AS events and limited event types. Second, they focused on AS detection tools that operate on the isoform level^6–11^, splice-aware mapping tools^12,13^, and differential splicing analysis tools^14–16^. However, AS detection tools that operate on the event level are more precise. Finally, benchmark studies are seldomly updated.

We used as ground truth simulated RNA-Seq data sets that were genome-wide and contained predefined number and distribution of AS events. To perform a standardised benchmark, we created a modular pipeline called DICAST (Docker Integration for Comparison of AS tools). It uses Docker to handle the installation process and Snakemake to handle AS analysis and the benchmark workflow. Its modularity allows for adding new tools in the future and for quickly updating the benchmark.

While our main focus was to benchmark AS event detection tools, we started from splice-aware mapping to provide the best possible input to those tools. Thus, we evaluated 11 splice-aware mapping tools in addition to eight AS detection tools. We demonstrate the benefits and limitations of AS event detection tools and suggest a putatively optimal strategy for comprehensive AS analysis.

## Results

### Tools for benchmark

We collected splice-aware mapping and AS event detection tools released between 2010 and 2021 based on a Pubmed and Google Scholar search with the following inclusion criteria. A tool should be (i) available, (ii) documented, (iii) under an open-source license, (iv) available as stand-alone software (web-services can usually not process a large amount of data), (v) used in project(s) other than described in the tool publication, (vi) able to process widely used file formats such as fasta/fastq, gtf/gff3, and bam/sam. Since we controlled the number of AS events in a custom genome annotation, a tool should be (vii) able to work with those as well. The final list of tools contains 11 splice-aware mapping tools and 8 AS detection tools (Table 1).

**Table 1.**
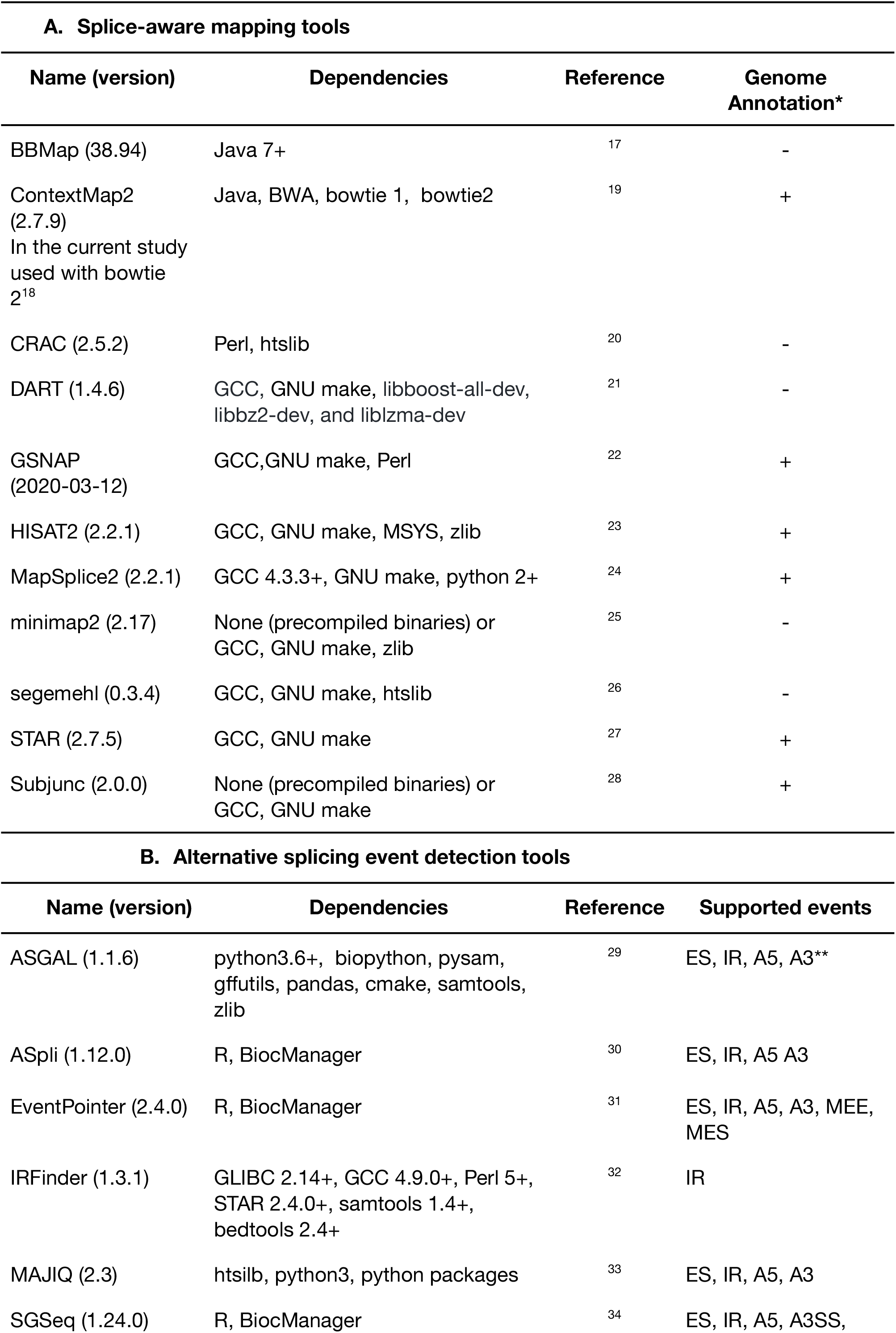
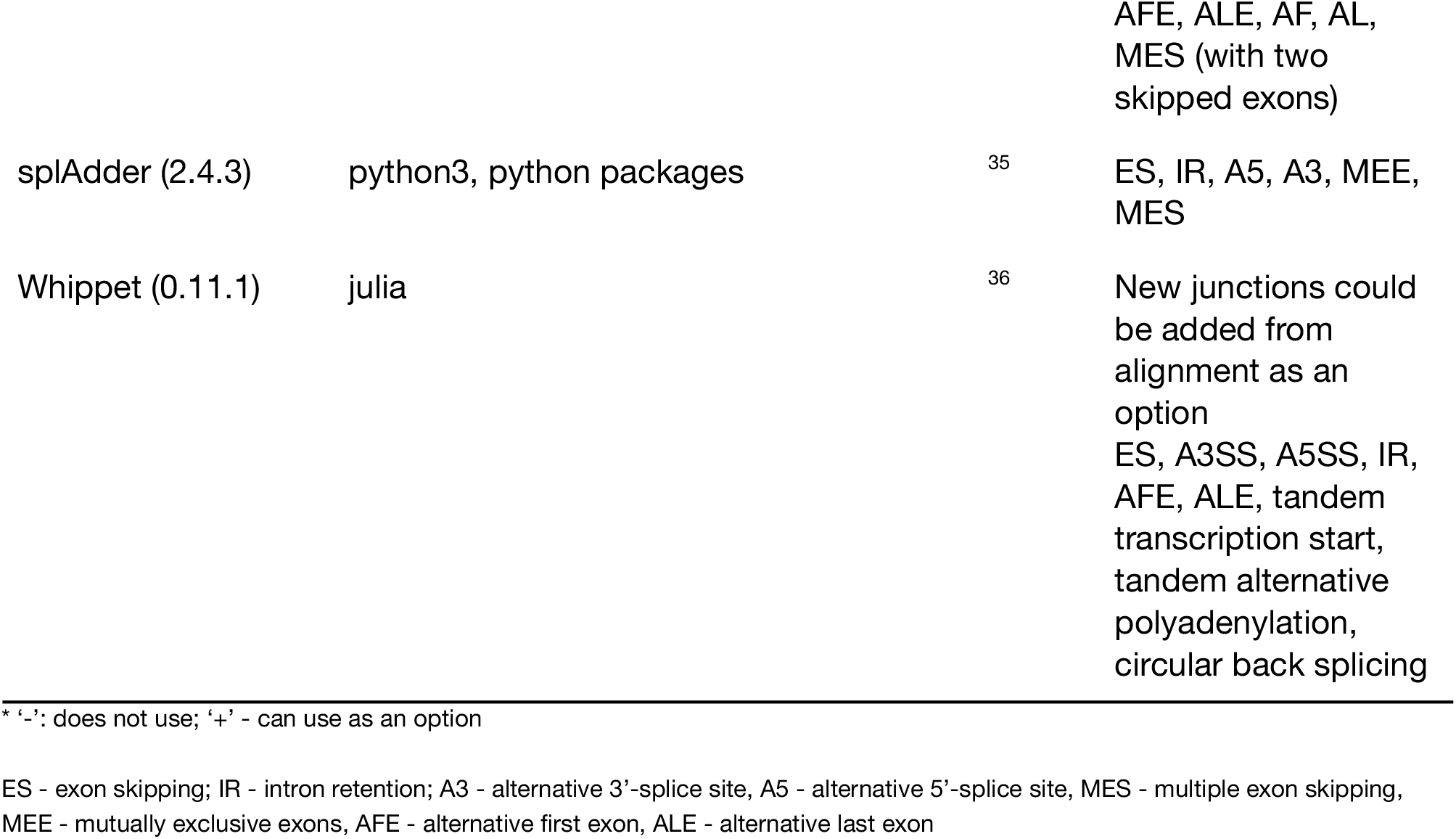
The list of AS analysis tools chosen for benchmark

### Alternative splicing analysis and benchmark pipeline

#### A unified output format for AS event detection

The main challenge for comparing AS event detection tools is the lack of standard output format. For an exon skipping event, ASGAL reports the coordinates of neighboring exons; ASpli reports the coordinates of a skipped exon itself; MAJIQ reports the coordinates of a neighboring exon and its junctions. We propose a unified format for all AS event types that reports the coordinates of (i) skipped exons for exon skipping, multiple exon skipping, mutually exclusive exons; (ii) a retained intron for intron retention; (iii) an alternative part of an exon for alternative splice sites (Supplementary Figure 1).

Additionally, for each event, the unified output format contains the gene name, chromosome, strand as well as a unique ID.

#### DICAST: Docker Integrated Comparison of Alternative Splicing Tools

The next necessity for the standardization of the AS analysis and benchmark process is a unified pipeline. DICAST handles (i) the installation process for every tool using Docker; (ii) the execution of tools using Snakemake; (iii) the output format unification; (iv) the comparison of AS events detected by different tools.

The general workflow starts with short-read RNA-Seq data (optionally simulated by ASimulatoR^37^, Figure 1) that serves as input for splice-aware mapping tools. The resulting alignment files then serve as input for AS event detection tools. Next, DICAST converts the output files of the AS event detection tools to the unified format described above, compares detected events across the tools, and reports the results of the comparison. A user could run all or separate steps of the workflow with their RNA-Seq reads, alignment, and genome annotation files.

**Figure 1.**
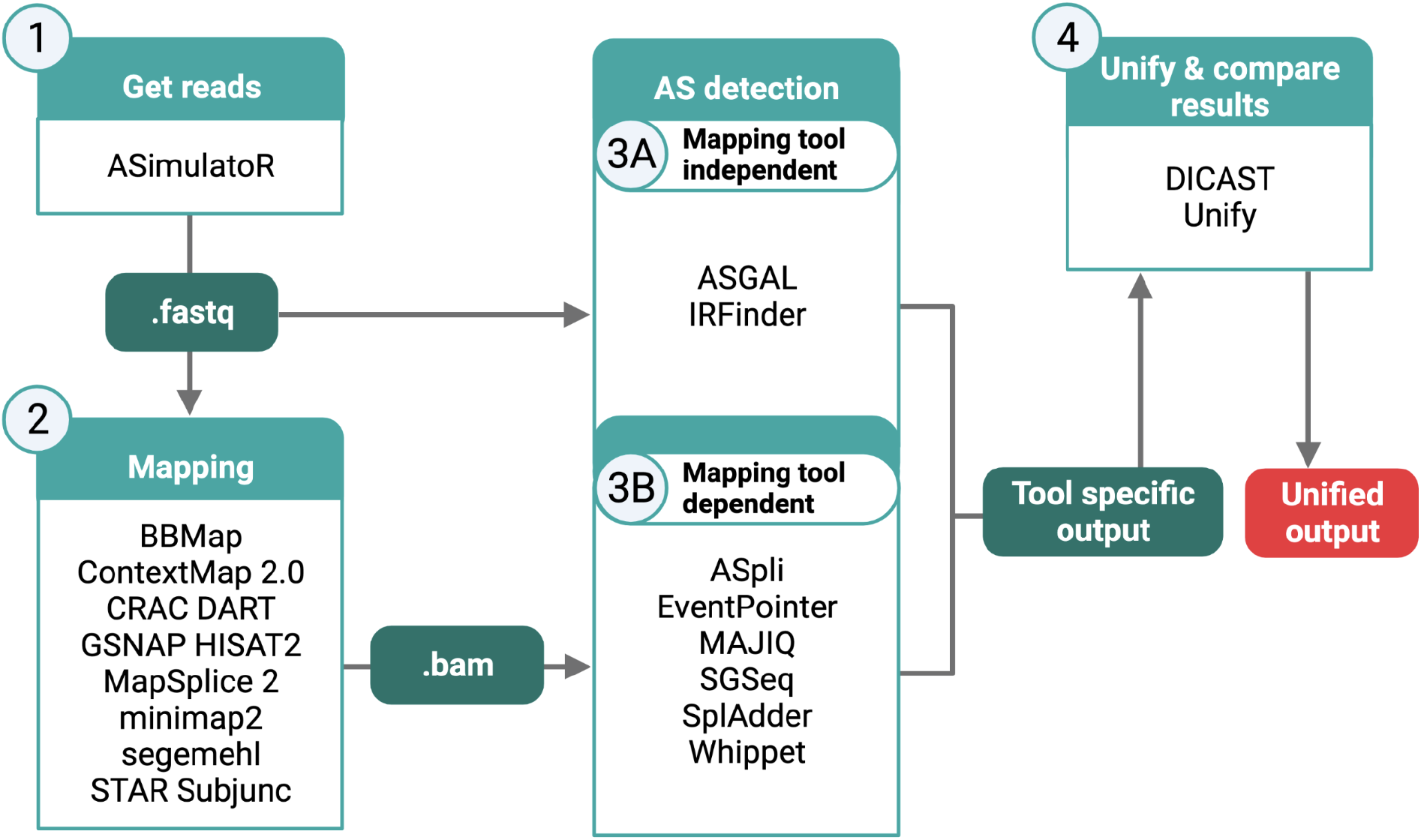
DICAST framework: 1) simulated (ASimulatoR) or user-provided fastq files 2) bam files could be generated by any of 11 supported splice-aware mapping tools; 3) AS events detected by any of 8 AS event detection tools based on files generated in the previous steps; 4) the output files of the AS detection tools are unified by DICAST. Created with BioRender.com

#### Benchmark workflow

For the benchmark, we simulated RNA-Seq data sets with 200 million paired-end reads (read length 76 base pairs) using ASimulatoR^37^. We tuned the complexity of the simulated data sets changing AS events distribution, combinations, and sequencing error rate (Table 2). The simulated data sets S1-S4 have an equal proportion of all event types while S5 mimics proportions from a biological data set.

**Table 2.** Characteristics of the simulated data sets

| Simulated Dataset | Events per transcript | Transcripts per gene | Events per exon | Event types |  |  |  |  |  |  | Sequencing error rate |
| --- | --- | --- | --- | --- | --- | --- | --- | --- | --- | --- | --- |
|  |  |  |  | ES | IR | A5 | A3 | MES | ALE | AFE |  |
| S1 | 1 | 1 | 1 | x | x | x | x | x | x | x | 0% |
| S2 | 1 | 1 | 1 | x | x | x | x | x | x | x | 0.1% |
| S3 | 2 | $\geq 1$ | 1 | x | x | x | x | x | x | x | 0.1% |
| S4 | 2 | $\geq 1$ | $\geq 1$ | x | x | x | x | x | x | x | 0.1% |
| S5 | 1-4 | $\geq 1$ | $\geq 1$ | x | x | x | x | | | | 0.1% |

Supplementary Figure 2 illustrates the workflow for benchmarking AS analysis tools. The simulated RNA-Seq data sets S1-S4 (Table 2) have an equal proportion of all types of AS events. We mapped these data sets to the human genome using 11 splice-aware mapping tools and calculated the proportion of unmapped reads relative to the number of all simulated reads (‘fraction of unmapped reads) and the proportion of correctly mapped reads and junctions relative to the number of all mapped reads (‘precision’). We chose the mapping tool with the best precision/fraction of unmapped reads values and used the resulting alignments as input for 8 AS detection tools. We then calculated the proportion of correctly found events relative to the number of events in the simulated set (‘recall’) and relative to the number of events found by the tool (‘precision’).

We also aimed to evaluate AS analysis tools in a biologically relevant setting but the real RNA-Seq data sets lack the genome-wide ground truth. We addressed this challenge and simulated a data set S5 with realistic event proportions, which were obtained by analyzing 117 RNA-Seq samples from the SHIP cohort (the Study of Health in Pomerania)^38^.

### Benchmark results: Splice-aware mapping tools

Splice-aware mapping tools differ in the mapping approach and their use of genome annotations. Most tools use variations of the seed-and-extend algorithm and start with aligning parts of a read (seeds). ContextMap and MapSplice2 first align reads end-to-end and use the seed-and-extend algorithm for reads that could not be aligned in the first step. Some tools can use genome annotations for additional information (e.g., splice sites) (Table 1), while others (e.g. BBmap) do not need it.

We provided a genome annotation where possible and used the best mode for AS analysis (e.g., a 2-pass mode for STAR). We downsampled the input files to 10 million reads uniformly at random to reduce the analysis time.

The mapping approach has a limited effect on the performance of the tool. ContextMap and MapSplice2 show comparable performance metrics as the tools with the best performance such as STAR and HISAT2 (Figure 2, Supplementary Figure 3). Tools that use genome annotation generally show better performance. Surprisingly, the complexity of the data sets only marginally affects the performance of the tools. Only DART suffers from a decrease in precision with an increasing sequencing error rate.

**Figure 2.**
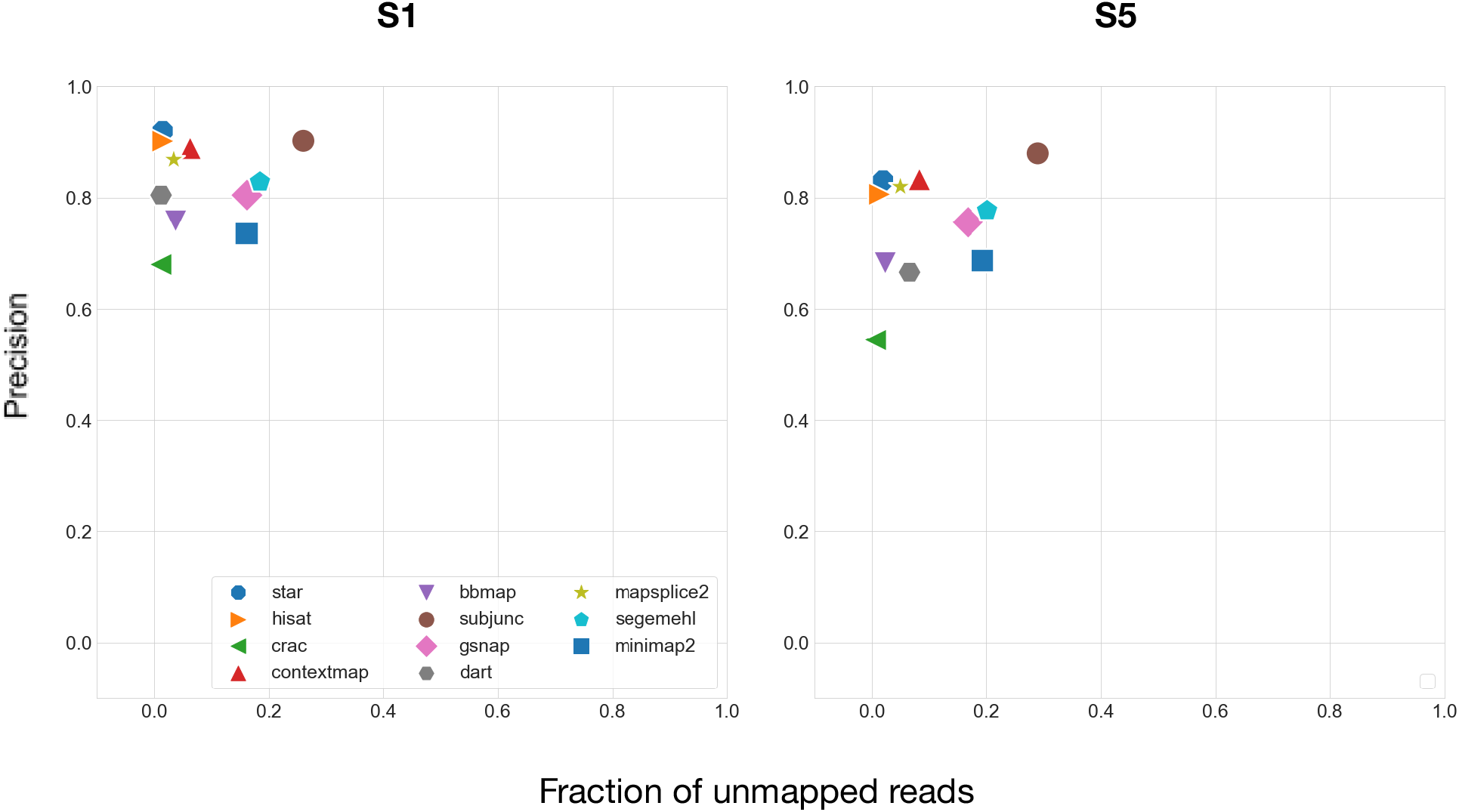
The plots for precision and fraction of unmapped reads plots for splice-aware mapping tools for the data sets S1 and S5. The results for S2-S4 are presented in Supplementary Figure 3.

We chose STAR for further analysis, as it demonstrated the best performance. We downsampled data sets to 200, 100, and 50 million reads uniformly at random to investigate the effect of sequencing depth on performance.

#### Alternative splicing detection tools

AS event detection tools detect events from genome annotation (ASpli, SGSeq (annotated transcripts), IRFinder), RNA-Seq alignment (EventPointer, SGSeq (*de novo*), MAJIQ), or both, i.e. by augmenting the genome annotation through alignment (ASGAL, splAdder, Whippet).

We excluded ASGAL because of the long run time (a genome-wide analysis of 50 million reads took around 3 days). As most tools (Table 1) do not detect mutually exclusive exons and multiple exon skipping, we treated such events as exon skipping events.

Figure 3A shows results for data sets S1 and S5. As for S1-S4 (Figure 3A, Supplementary Figure 4), the ranking of the tools differs slightly. Whippet and SGSeq_Anno (annotated transcripts) show the best performance. EventPointer and SGSeq_denovo (*de novo*) demonstrate poor performance. For all tools, recall depends on the sequencing depth, while precision does not.

**Figure 3.**
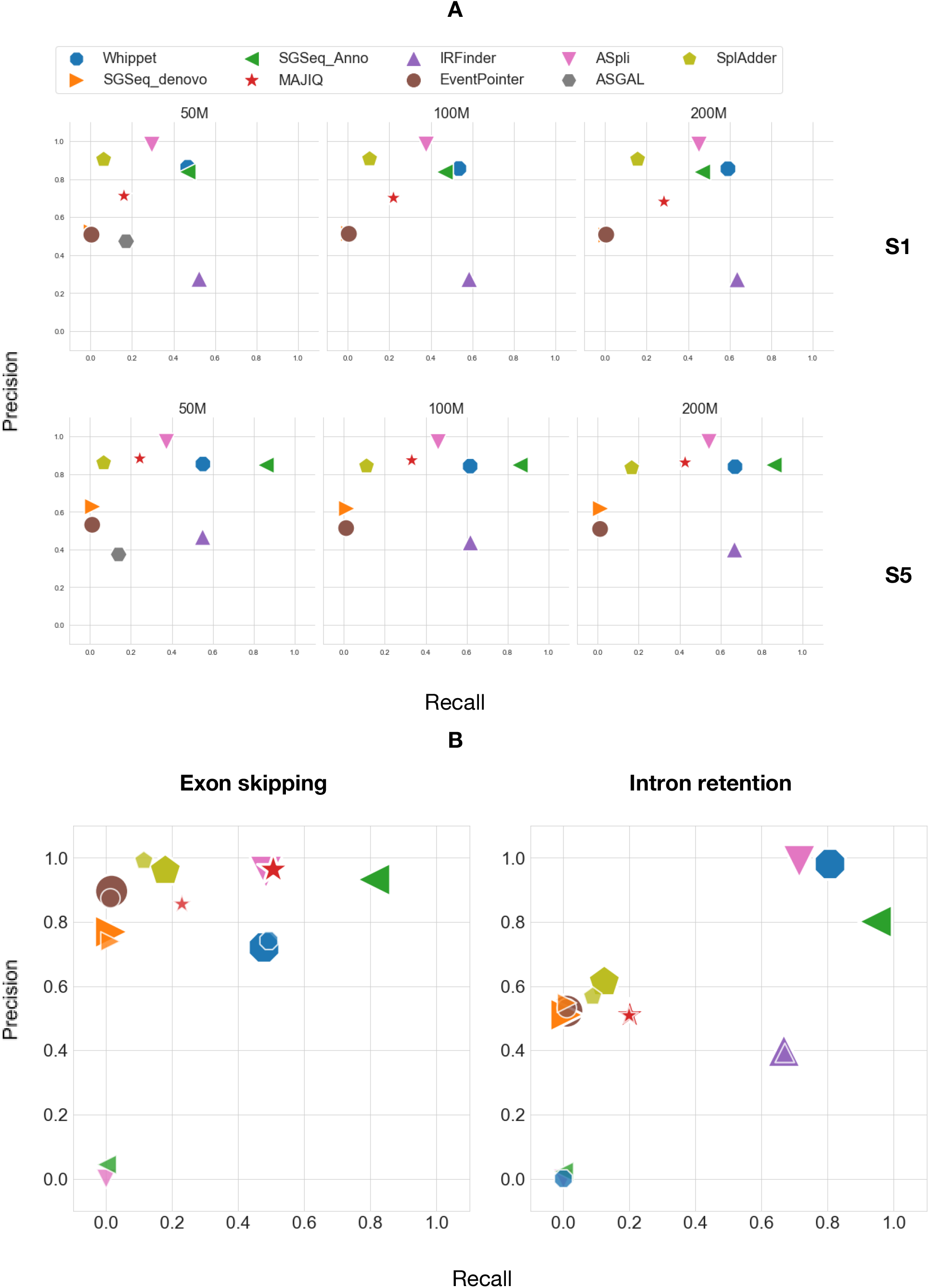
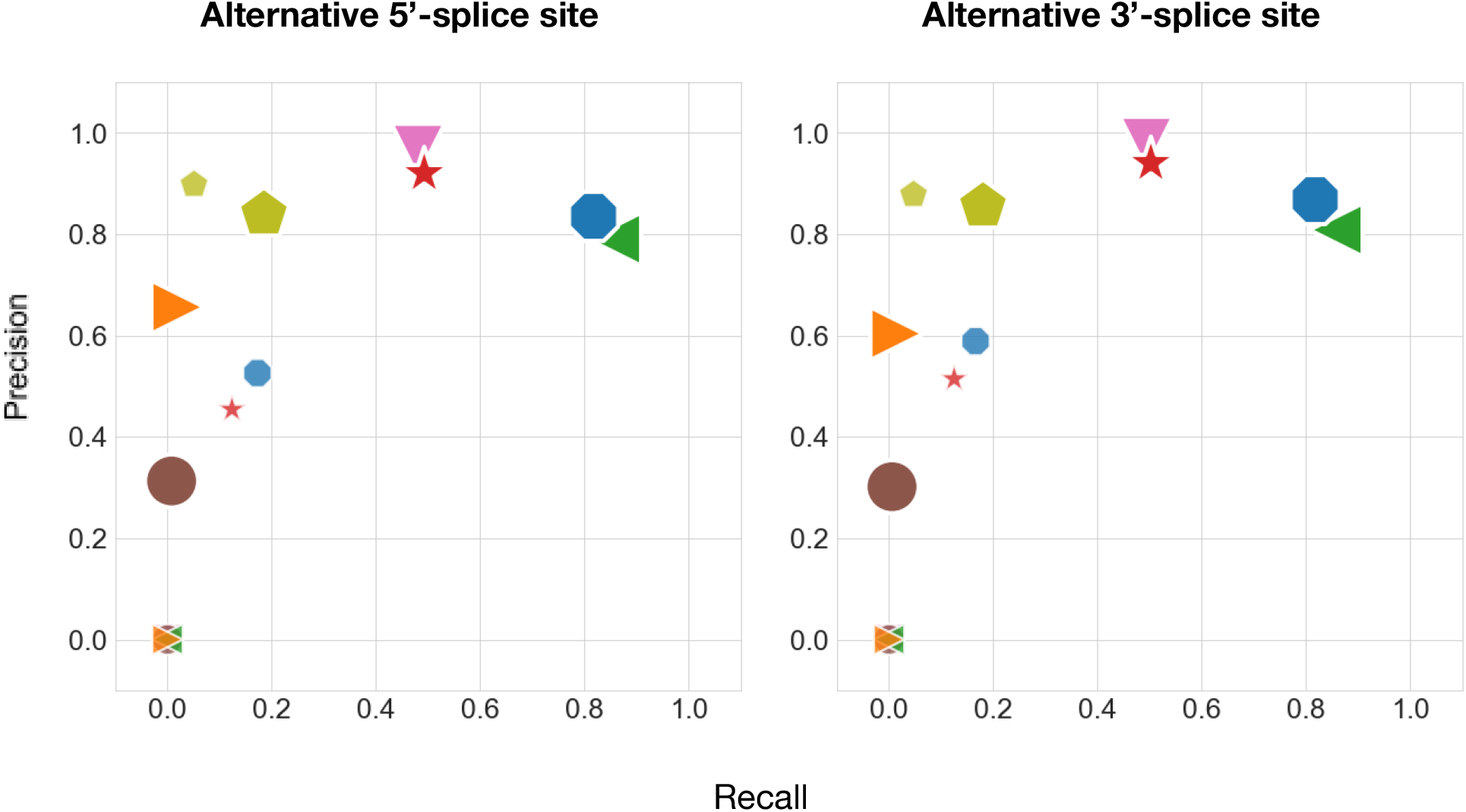
A. Precision/recall plots for AS event detection tools for data sets S1 and S5. B. Tool performance on S5 (bigger markers) and S5-tr (smaller markers) sets by AS event type.

MAJIQ was used to derive the proportion of AS event types and combinations from the SHIP cohort (Supplementary Figure 5). We used the proportions as a parameter for ASimulatoR to simulate the biologically-inspired set S5. Note that only the proportion values, but not the detected AS events were used, so MAJIQ and the other tools remained unaware of exact AS events in S5. We used MAJIQ here, since it does not depend on genome annotation and can extract events directly from alignments. Additionally, we truncated the resulting genome annotation file to retain only one transcript per gene and investigated the ability of tools to detect events *de novo*. We denote this data set as S5-tr.

For MAJIQ and SGSeq (annotated transcripts), we observed an increase in precision and/or recall in the data set S5 (Figure 3A), likely because S5 contains single but not multiple exon skipping events (e.g., mutually exclusive exons). This observation demonstrates that these tools might underestimate AS events that involve multiple exons.

The tested tools differ significantly in their abilities to detect *de novo* events based on truncated genome annotation. ASpli and SGSeq (annotated transcripts) detected almost no novel events. For other tools, we compared precision and recall for S5 and S5-tr for each AS event type individually (Figure 3B). With the full annotation, intron retention was the most difficult type of event to recover. Whippet did not find any *de novo* intron retention events. The recovery rate of alternative splice sites also dropped for all tools. The most dramatic changes we observed for MAJIQ (precision decreased from around 0.9 to 0.5; recall decreased from 0.5 to 0.1) and Whippet (precision decreased from around 0.8 to 0.5; recall decreased from 0.8 to 0.2)

Hypothesizing that combining several tools might improve performance, we considered combinations of two or more tools (OR) as well as their overlap (AND) for the best-performing individual tools IRFinder, MAJIQ, splAdder, and Whippet (Supplementary Figure 6).

Combining the results of two or more tools (OR) leads to a loss in precision without meaningfully increasing recall. Using the overlapping results (AND), there is a gain in precision but a loss in recall. The latter approach might be appropriate if high-confident AS events are needed concerning the possibility of low overlap between multiple tools.

In summary, for the comprehensive analysis of AS events from short-read RNA-Seq data, we suggest the following approach: (i) detect known events using ASpli, Whippet, or SGSeq (annotated transcripts); (ii) detect intron retention *de novo* using IRFinder; (iii) detect other *de novo* events balancing between tools with better precision (splAdder) and tools with better recall (Whippet and MAJIQ).

#### Run time

Using the data set S5, we estimated the run time of the tools tested on Intel® Xeon® Gold 6148 Processor with 27.5M Cache, 2.40 GHz (Figure 4). For the mapping tools, we used the downsampled data set with 10 million reads from the benchmark. Most tools perform indexing and mapping within an hour. ContextMap is the slowest tool with a run time of around 11 hours. minimap2 and BBmap are the fastest tools with a run time of only several minutes. We also estimated the run time of AS detection tools depending on the sequencing depth (Figure 4). The run time of MAJIQ does not depend on the sequencing depth and is in the range of minutes. Most other tools run within several hours, with splAdder taking up to 10 hours for 200 million reads. ASGAL was only evaluated with 50 million reads, already running longer than 72 hours to complete.

**Figure 4.**
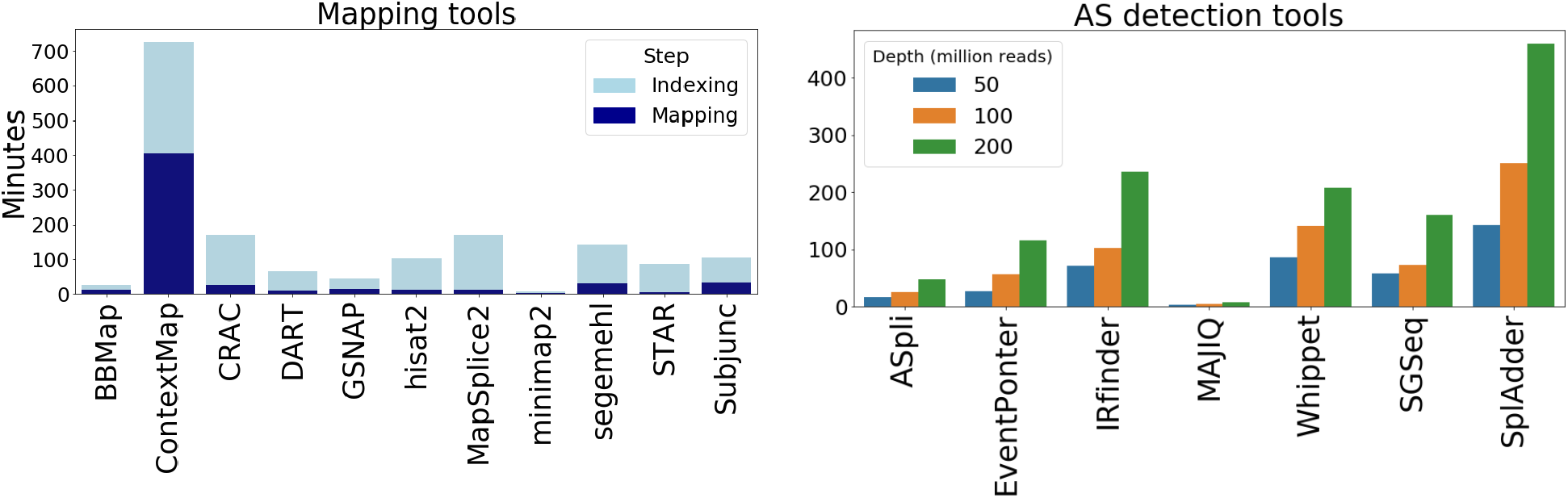
The run time in minutes of the splice-aware mapping tools and AS event detection tools.

## Conclusion

We investigated the performance of AS event detection tools in a comprehensive benchmark. We started with splice-aware mapping tools to obtain the best possible input for event detection. Among splice-aware mapping tools, STAR and HISAT2 present the best balance between the precision value, fraction of unmapped reads values, and the runtime, which is in concordance with previous findings^12^. For organisms without genome annotation, BBmap is promising as it does not need a genome annotation, has relatively good precision value, low fraction of unmapped reads, and short run time.

Concerning AS event detection, we still find much room for improvement since tools with high recall values (Whippet, SGSeq (annotated transcripts), ASpli) can not detect events *de novo*. Vice versa, tools that can identify events *de novo* demonstrate poor recall values. We suggest using a combination of existing tools for a comprehensive AS analysis on the event level using short-read RNA-Seq data.

While we aimed for a comprehensive, objective, and standardized benchmark, we faced some limitations. First, the simulated data sets might not reach the same level of complexity as real biological data sets. We mitigated this by mimicking the proportions of AS event types as observed in the SHIP cohort. Second, for run time reasons, we refrained from a tool-specific parameter tuning and relied on the default parameters, which should ideally lead to optimal results in typical settings. Additionally, we did not want to favor tools with higher flexibility simply due to an increased number of parameters to tune. Finally, we did not explore the performance of the machine and deep learning-based tools (e.g. DARTS^39^, IRFinder-S^40^). The simulated AS events were added randomly and do not account for, e.g., regulatory sequences that are used in machine learning approaches.

Many challenges in AS analysis have yet to be addressed, including general difficulties for tool development such as the lack of efficient parallelization and substantial run time. We found that the standard format for aligned reads - bam/sam/cram - differs between tools and might not be compatible with some AS event detection tools. This fact makes many splice-aware alignment tools useless for AS analysis. While DICAST introduces a unifying standard for AS event reporting, AS event detection tools utilize inherently different approaches and lead to inconsistent results. To mitigate this, DICAST allows users to execute any combination of tools and facilitates adding newly published tools. By standardizing the output of AS event detection tools, DICAST significantly simplifies downstream analysis. In summary, DICAST offers a unified interface for existing methods and boosts method development by offering an easily extensible framework for benchmarking of existing and novel AS analysis approaches.

## Methods

### DICAST

DICAST uses Docker for full reproducibility and to simplify deployment^41,42^. Each docker image in DICAST can be used individually or as part of the benchmark/AS analysis pipeline. DICAST orchestrates the pipeline using Snakemake^43^. The DICAST source code and documentation are available at https://github.com/CGAT-Group/DICAST and https://dicast.readthedocs.io/en/master/contents.html, respectively.

### Benchmark

RNA-Seq datasets were simulated using the R package ASimulatoR^37^ with a sequencing depth of 200 million reads based on the human genome hg38 and the Ensembl genome annotation, version 99^44^. We limited the simulation to chromosomes 1-22, X, Y, and MT.

For each simulated dataset, ASimulatoR estimated how many genes have enough exons to create AS events and can, hence, be used for the simulation. For the simulated set S3, this number was the lowest - 37648. We used this number of genes as a parameter for all simulated datasets to preserve the read coverage level.

We further used seqtk (available at https://github.com/lh3/seqtk) to downsample fastq files uniformly at random to 50 and 100 million reads for benchmarking AS detection tools and 10 million reads for the mapping tools benchmark.

Precision and fraction of unmapped reads for splice-aware mapping tools were calculated based on the resulting alignment file and the correct coordinates provided by ASimulatoR. Precision and recall for AS event detection tools were calculated based on the unified output of DICAST and the description of correct events provided by ASimulatoR.

### SHIP cohort

The Study of Health in Pomerania (SHIP) is a longitudinal population-based cohort study located in the area of West Pomerania (Northeast Germany). For RNA-Seq analysis, total RNA was extracted from whole blood with a mean RNA integrity of 8.5. Based on 500ng total RNA per sample, a library was prepared using the TruSeq Stranded mRNA kits (A and B) with 24 barcodes and 117 samples were sequenced on Illumina HiSeq 4000, 2x 101bp paired-end reads with a sequencing depth ∼ 40Mio clusters per sample.

Written informed consent was obtained from SHIP-TREND study participants, and all protocols were approved by the institutional ethical review committee in adherence with the Declaration of Helsinki.

We used these 117 RNA-Seq samples from the SHIP-TREND cohort^38^ in this study. They were mapped to the human genome hg38 with STAR using the Ensembl genome annotation, version 99, and analyzed further with MAJIQ^33^. Custom scripts in Python 3 were used to obtain the number of AS events and genes with AS.

## Data availability

Access to the SHIP data for research purposes may be requested at https://www.fvcm.med.uni-greifswald.de/dd_service/data_use_intro.php.

## Authors contribution

Planning and concept of study: TK, ML; Data analysis: AF, OT, AD, TF, JK, CTL; DICAST development: AF, TF, FR, AD, JK, ZL, CTL. Manuscript writing and revision: all authors.

## Acknowledgments

NGS analyses were carried out at the MDC (Berlin) in the frame of the “*OMICs initiative”* of the DZHK (German Centre for Cardiovascular Research).

The authors thank Dr. Sabine Ameling for her help with the description of the experimental procedures. We also thank Alexander Gress and Lena Maria Hackl for testing the DICAST pipeline.

## Supplementary Materials

**Supplementary Figure 1.**
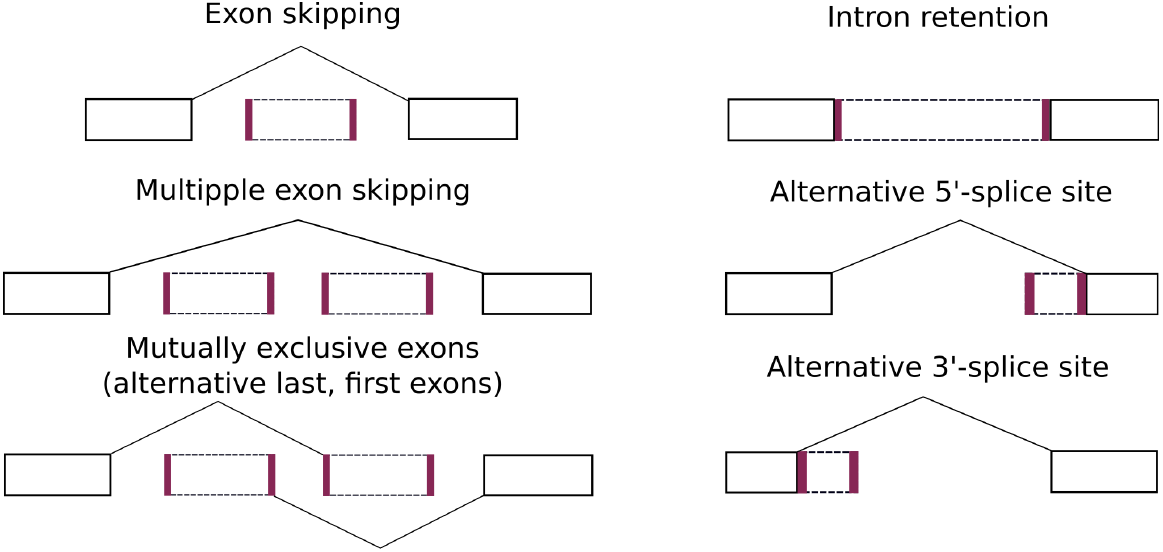
The coordinates reported in the unified format output of DICAST or AS events are shown as bold purple lines in the corresponding splice graphs.

**Supplementary Figure 2.**
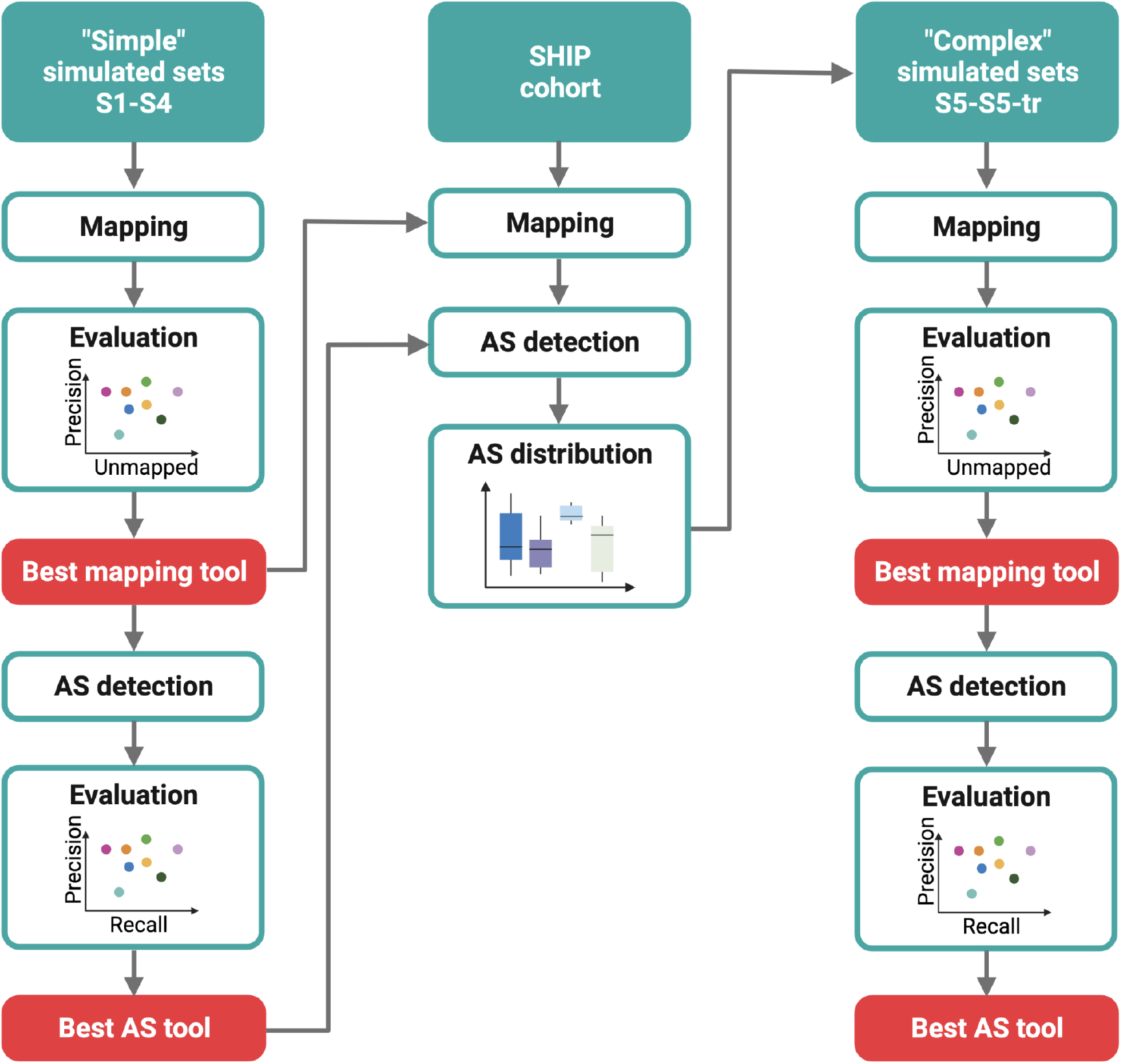
The workflow used to benchmark AS analysis tools. Created with BioRender.com.

**Supplementary Figure 3.**
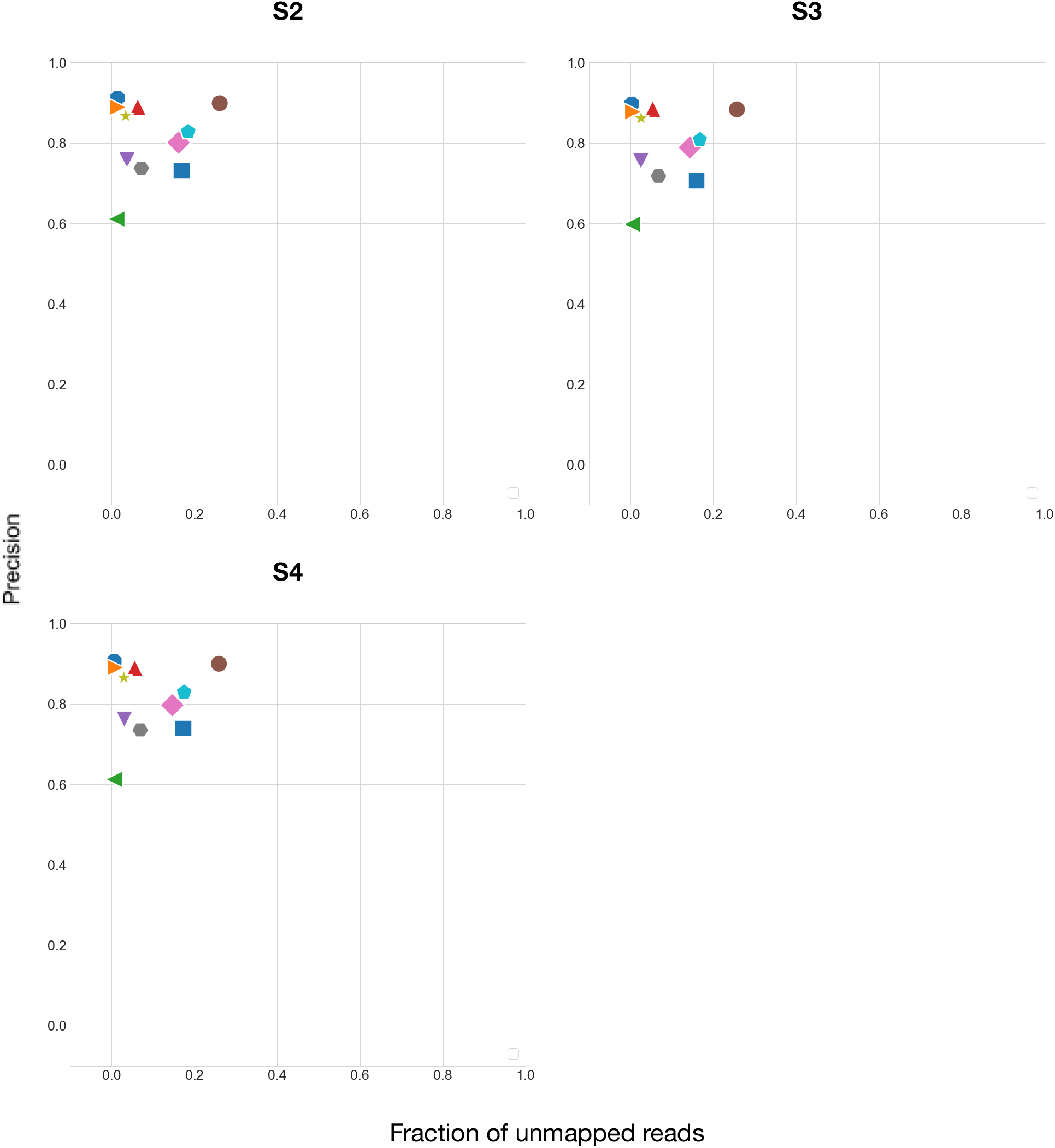
The plots for precision and fraction of unmapped reads for splice-aware mapping tools for the S2-S4 simulated sets.

**Supplementary Figure 4.**
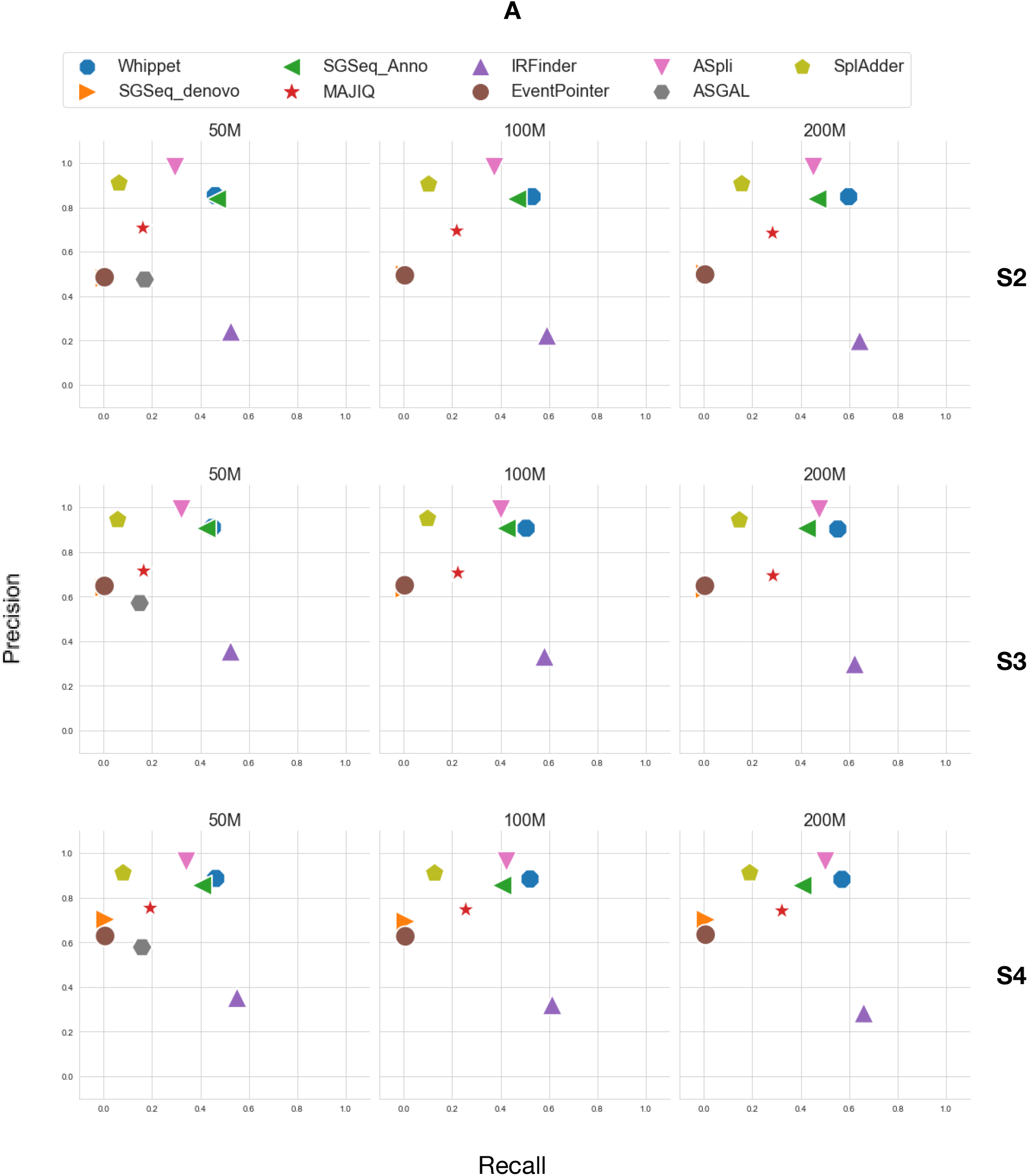
Precision/recall plots for AS event detection tools for the S2-S4 simulated sets

**Supplementary Figure 5.**
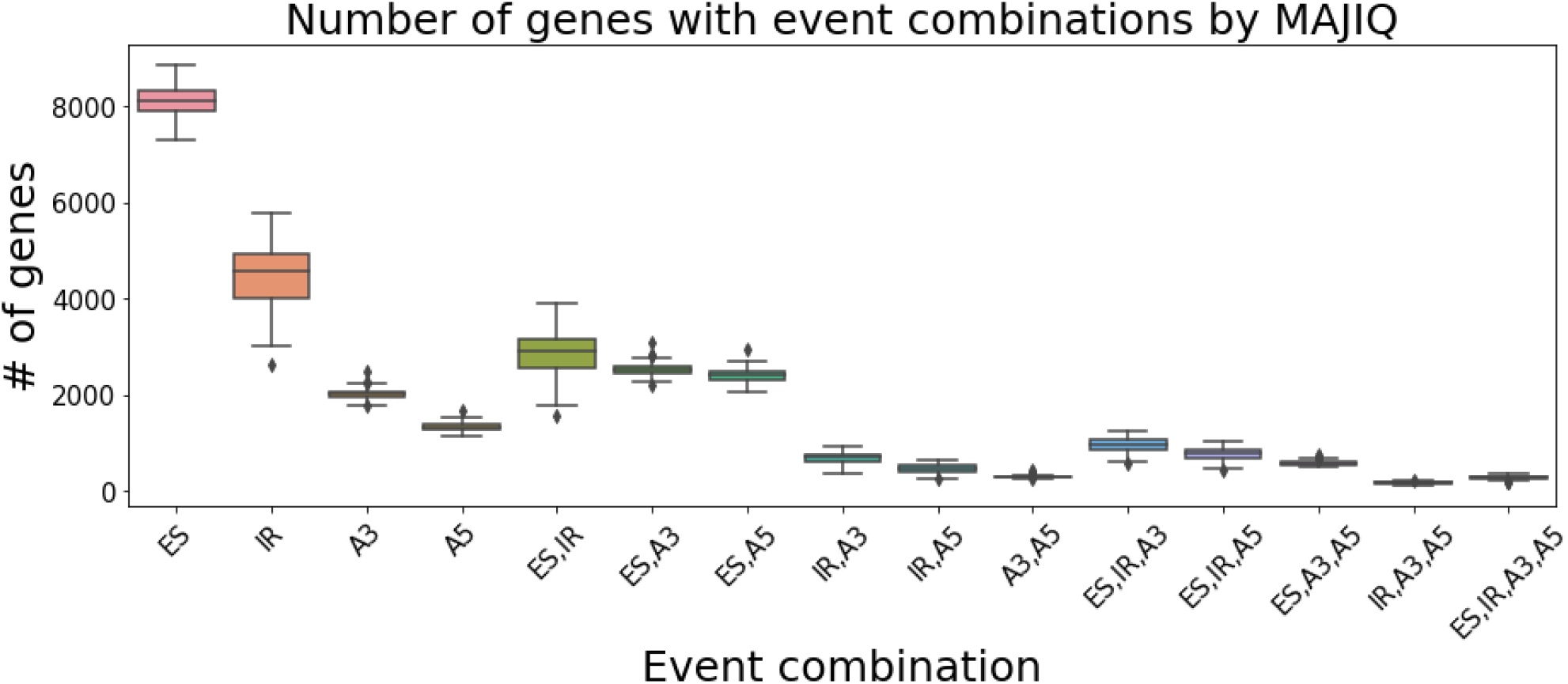
Number of genes with an AS event found by MAJIQ in the SHIP cohort. ES: Exon skipping, IR: Intron retention, A3: Alternative 3’-splice site, A5: Alternative 5’-splice site.

**Supplementary Figure 6.**
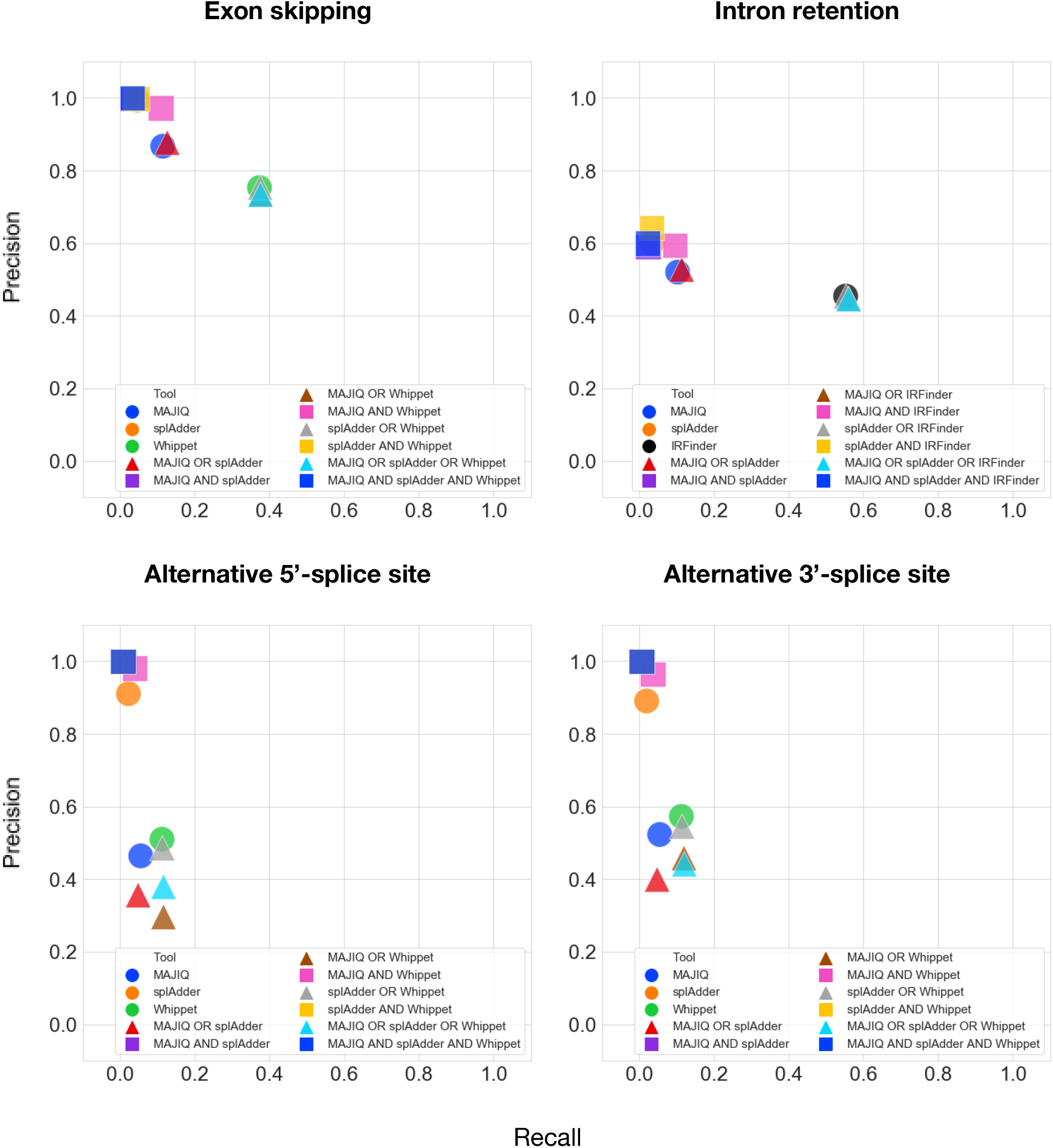
Precision/recall plots for the results of individual and combined AS event detection tools divided by AS event type.

## References

1. Pan, Q., Shai, O., Lee, L. J., Frey, B. J. & Blencowe, B. J. Deep surveying of alternative splicing complexity in the human transcriptome by high-throughput sequencing. Nature Genetics vol. 40 1413–1415 (2008).

2. Wang, E. T. et al. Alternative isoform regulation in human tissue transcriptomes. Nature 456, 470–476 (2008).

3. Bonnal, S. C., López-Oreja, I. & Valcárcel, J. Roles and mechanisms of alternative splicing in cancer - implications for care. Nat. Rev. Clin. Oncol. 17, 457–474 (2020).

4. Nikonova, E., Kao, S.-Y. & Spletter, M. L. Contributions of alternative splicing to muscle type development and function. Semin. Cell Dev. Biol. 104, 65–80 (2020).

5. Zheng, S. Alternative splicing programming of axon formation. Wiley Interdiscip. Rev. RNA 11, e1585 (2020).

6. Zhang, C., Zhang, B., Lin, L.-L. & Zhao, S. Evaluation and comparison of computational tools for RNA-seq isoform quantification. BMC Genomics 18, 583 (2017).

7. Jin, H., Wan, Y.-W. & Liu, Z. Comprehensive evaluation of RNA-seq quantification methods for linearity. BMC Bioinformatics 18, 117 (2017).

8. Dapas, M., Kandpal, M., Bi, Y. & Davuluri, R. V. Comparative evaluation of isoform-level gene expression estimation algorithms for RNA-seq and exon-array platforms. Brief. Bioinform. 18, 260–269 (2017).

9. Hayer, K. E., Pizarro, A., Lahens, N. F., Hogenesch, J. B. & Grant, G. R. Benchmark analysis of algorithms for determining and quantifying full-length mRNA splice forms from RNA-seq data. Bioinformatics 31, 3938–3945 (2015).

10. Leshkowitz, D. et al. Using Synthetic Mouse Spike-In Transcripts to Evaluate RNA-Seq Analysis Tools. PLoS One 11, e0153782 (2016).

11. Chandramohan, R., Wu, P.-Y., Phan, J. H. & Wang, M. D. Benchmarking RNA-Seq quantification tools. Conf. Proc. IEEE Eng. Med. Biol. Soc. 2013, 647–650 (2013).

12. Baruzzo, G. et al. Simulation-based comprehensive benchmarking of RNA-seq aligners. Nat. Methods 14, 135–139 (2017).

13. Engström, P. G. et al. Systematic evaluation of spliced alignment programs for RNA-seq data. Nat. Methods 10, 1185–1191 (2013).

14. Mehmood, A. et al. Systematic evaluation of differential splicing tools for RNA-seq studies. Brief. Bioinform. 21, 2052–2065 (2020).

15. Liu, R., Loraine, A. E. & Dickerson, J. A. Comparisons of computational methods for differential alternative splicing detection using RNA-seq in plant systems. BMC Bioinformatics 15, 364 (2014).

16. Merino, G. A., Conesa, A. & Fernández, E. A. A benchmarking of workflows for detecting differential splicing and differential expression at isoform level in human RNA-seq studies. Brief. Bioinform. 20, 471–481 (2019).

17. Marić, J. Long Read RNA-seq Mapper. (2015).

18. Langmead, B. & Salzberg, S. L. Fast gapped-read alignment with Bowtie 2. Nat. Methods 9, 357–359 (2012).

19. Bonfert, T., Kirner, E., Csaba, G., Zimmer, R. & Friedel, C. C. ContextMap 2: fast and accurate context-based RNA-seq mapping. BMC Bioinformatics 16, 122 (2015).

20. Philippe, N., Salson, M., Commes, T. & Rivals, E. CRAC: an integrated approach to the analysis of RNA-seq reads. Genome Biol. 14, R30 (2013).

21. Lin, H.-N. & Hsu, W.-L. DART: a fast and accurate RNA-seq mapper with a partitioning strategy. Bioinformatics 34, 190–197 (2018).

22. Wu, T. D. & Nacu, S. Fast and SNP-tolerant detection of complex variants and splicing in short reads. Bioinformatics 26, 873–881 (2010).

23. Kim, D., Paggi, J. M., Park, C., Bennett, C. & Salzberg, S. L. Graph-based genome alignment and genotyping with HISAT2 and HISAT-genotype. Nat. Biotechnol. 37, 907–915 (2019).

24. Wang, K. et al. MapSplice: accurate mapping of RNA-seq reads for splice junction discovery. Nucleic Acids Res. 38, e178 (2010).

25. Li, H. Minimap2: pairwise alignment for nucleotide sequences. Bioinformatics 34, 3094–3100 (2018).

26. Hoffmann, S. et al. Fast mapping of short sequences with mismatches, insertions and deletions using index structures. PLoS Comput. Biol. 5, e1000502 (2009).

27. Dobin, A. et al. STAR: ultrafast universal RNA-seq aligner. Bioinformatics 29, 15–21 (2013).

28. Appleby, L. & Araya, R. Postgraduate training in psychiatry 1977-87: disturbing trends in the pattern of international collaboration. Med. Educ. 24, 290–297 (1990).

29. Denti, L. et al. ASGAL: aligning RNA-Seq data to a splicing graph to detect novel alternative splicing events. BMC Bioinformatics 19, 444 (2018).

30. Estefania, M., Andres, R., Javier, I., Marcelo, Y. & Ariel, C. ASpli: Integrative analysis of splicing landscapes through RNA-Seq assays. Bioinformatics (2021) doi:10.1093/bioinformatics/btab141.

31. Romero, J. P. et al. EventPointer: an effective identification of alternative splicing events using junction arrays. BMC Genomics 17, 467 (2016).

32. Middleton, R. et al. IRFinder: assessing the impact of intron retention on mammalian gene expression. Genome Biol. 18, 51 (2017).

33. Vaquero-Garcia, J. et al. A new view of transcriptome complexity and regulation through the lens of local splicing variations. Elife 5, e11752 (2016).

34. Goldstein, L. D. et al. Prediction and Quantification of Splice Events from RNA-Seq Data. PLoS One 11, e0156132 (2016).

35. Kahles, A., Ong, C. S., Zhong, Y. & Rätsch, G. SplAdder: identification, quantification and testing of alternative splicing events from RNA-Seq data. Bioinformatics 32, 1840–1847 (2016).

36. Sterne-Weiler, T., Weatheritt, R. J., Best, A. J., Ha, K. C. H. & Blencowe, B. J. Efficient and Accurate Quantitative Profiling of Alternative Splicing Patterns of Any Complexity on a Laptop. Mol. Cell 72, 187–200.e6 (2018).

37. Manz, Q. et al. ASimulatoR: splice-aware RNA-Seq data simulation. Bioinformatics (2021) doi:10.1093/bioinformatics/btab142.

38. Völzke, H. [Study of Health in Pomerania (SHIP). Concept, design and selected results]. Bundesgesundheitsblatt Gesundheitsforschung Gesundheitsschutz 55, 790–794 (2012).

39. Zhang, Z. et al. Deep-learning augmented RNA-seq analysis of transcript splicing. Nat. Methods 16, 307–310 (2019).

40. Lorenzi, C. et al. IRFinder-S: a comprehensive suite to discover and explore intron retention. Genome Biol. 22, 307 (2021).

41. Merkel, D. Docker: lightweight Linux containers for consistent development and deployment. Linux J. 2014, 2 (2014).

42. Nüst, D. et al. Ten simple rules for writing Dockerfiles for reproducible data science. PLoS Comput. Biol. 16, e1008316 (2020).

43. Mölder, F. et al. Sustainable data analysis with Snakemake. F1000Res. 10, (2021).

44. Yates, A. D. et al. Ensembl 2020. Nucleic Acids Res. 48, D682–D688 (2020).

